# Deep learning facilitates precise identification of disease-resistance genes in plants

**DOI:** 10.1101/2024.09.26.615248

**Authors:** Zhenya Liu, Xu Wang, Shuo Cao, Tingyue Lei, Yifu Chenzhu, Mengyan Zhang, Zhongqi Liu, Jianzhong Lu, Wenqi Ma, Bingxiong Su, Yiwen Wang, Yongfeng Zhou

**Author notes:** These authors contributed equally to this work (Zhenya Liu, Xu Wang, Shuo Cao).

## Abstract

The identification of plant disease-resistance genes is essential for understanding the plant immune system and accelerating crop breeding with disease resistance. Therefore, there is a pressing need for a method capable of accurately and comprehensively annotating resistance genes at a genome-wide scale. In this study, we propose Evolutionary Scale Modeling for LRR (ESM-LRR), a novel approach based on the deep protein language model to accurately identify LRR domains which are substantially variable structures in disease-resistance proteins. ESM-LRR achieved its highest F1 score of 0.80 on a test set using the 90% identity as the matching threshold. Building upon ESM-LRR, we developed the Plant Disease-Resistance Gene Predictor (R-Predictor), a framework designed to simultaneously annotate 15 diverse domain topologies, extensively covering recently characterised resistance genes across the entire genome. R-Predictor integrates four modules, each employing superior methods that outperform existing methods (F1 score of 0.89 for RLKs and 0.88 for NLRs), demonstrating its high accuracy and practicality in annotating disease-resistance genes (R genes). R-Predictor were also applied to identify R genes in rice, tomato, and grapevine. Furthermore, AlphaFold3 was employed to screen interactions between 1,116 protein pairs involving three NLRs identified exclusively by R-Predictor and 372 literature-reported plant-pathogen effectors, resulting in the identification of 15 putative NLR-effector complexes. Overall, this study presents a novel approach for advancing our understanding of plant immune mechanisms.

## Introduction

Plant diseases are responsible for an average loss of 20-30% in global crop yields^1, 2^. In 2019, approximately two million tons of agrichemicals were used in crop production, with 47.5% allocated to herbicides, 29.5% to insecticides, and 17.5% to fungicides^3^. Scientists are dedicated to mitigating these losses and reducing the reliance on agrichemicals by enhancing plant immunity. The plant immune system is conceptualized into a “zig-zag-zig” intellectual framework, which consists of pattern-triggered immunity (PTI), effector-triggered susceptibility (ETS), and effector-triggered immunity (ETI)^4^. The recognition of pathogen-, damage-, microbe-, or herbivore-associated molecular patterns (PAMPs, DAMPs, MAMPs, HAMPs) by pattern-recognition receptors (PRRs) on the plant cell surface initiates defense responses against invading pathogens, a process known as PTI^5^. PRRs include receptor-like kinases (RLKs) and receptor-like proteins (RLPs), the latter of which lack a protein kinase domain. To counteract or suppress PTI, pathogens have evolved to secrete specific effectors, resulting in effector-triggered susceptibility (ETS)^6^. However, plants have also evolved intracellular nucleotide-binding leucine-rich repeat receptors (NLRs) that can directly or indirectly recognize these effectors, thereby activating ETI^7, 8^. Consequently, the precise identification of disease-resistance genes (R genes) involved in the production of RLKs, RLPs and NLRs is essential for the enhancement of plant immunity.

Typically, R genes are not always polymorphic in natural plant populations^9^. Plant RLKs comprise a signal peptide, an ectodomain (ECD), a transmembrane domain and a protein kinase domain, whereas RLPs are essentially RLKs that lack the protein kinase domain^10^. The leucine-rich repeat (LRR)-RLKs and LRR-RLPs are the most abundant subfamilies of RLKs and RLPs, respectively, with the LRR domain serving as their ECD. The LRR domain is a widespread structural motif consisting of two or more LRR units, each comprising 20-30 amino acids, characterized by a sequence pattern rich in Leucine. The LRR unit typically repeats in tandem, forming a curved solenoid structure that facilitates protein-protein interactions^11^. Similar to RLKs and RLPs, plant NLRs are modular proteins generally composed of three domains: an N-terminal domain, a central NB-ARC domain and a C-terminal LRR domain. Plant NLRs can be divided into TIR-type NLRs (TNLs) and CC-type NLRs (CNLs), distinguished by the presence of either a TOLL/interleukin-1 receptor ^12^ domain or a coiled-coil (CC) domain at their N-terminal domain respectively. However, some R genes lack typical domain combinations or topologies, exhibiting replaced, missing or additional domains, such as CERK1^13^, EDR1^14^, CEBiP^13, 15^, NRG1^16^ and DAR5^17^, yet they still play crucial roles in plant immunity. Therefore, characterizing the domain topology of R genes is essential for identifying potentially overlooked resistance genes and mechanisms.

High-quality disease-resistance gene annotations have been established for the model species *Arabidopsis thaliana* through collaborative community efforts and meticulous manual curation^18^. However, with the rapid advancement of sequencing technologies, numerous reference genomes have been assembled both within and across species, creating an urgent demand for automated annotation methods for R genes in crops and their wild relatives. To date, several methods have been developed to accomplish this goal, including NLRtracker^19^, DeepLRR^20^, NLRexpress^21^, RLKdb^22^ and Resistify^23^. Nevertheless, these methods exhibit various limitations. For instance, RLKdb is restricted to identifying RLKs, while NLRtracker, NLRexpress and Resistify are confined to identifying NLRs. Although DeepLRR can identify RLKs, RLPs and NLRs simultaneously, it is constrained to typical domain structures. To overcome these individual shortcomings, different methods have been combined to achieve complementary results. However, no comprehensive benchmarking has been conducted to identify the optimal combination of methods for annotating the same type of receptors.

Furthermore, the LRR domain, a critical domain of RLKs, RLPs and NLRs exhibits substantial variability, which poses challenges for accurate identification using existing methods. To address this, we propose a novel approach called Evolutionary Scale Modeling for LRR (ESM-LRR), which leverages the deep protein language model ESM-1v^24^ to accurately characterize these variable LRR domains. The application of deep learning in bioinformatics has experienced rapid expansion and demonstrated exceptional potential in uncovering complex patterns within large-scale biological datasets^25, 26^. For instance, ResGitDR has been developed to predict chemotherapy agent responses based on somatic genome alterations^27^, while RNAErnie was designed for the annotation of RNA types^28^. Additionally, deep protein language models have been demonstrated to accurately estimate the effects of amino acid residue changes within protein sequences^29^. Therefore, the development of deep learning-based methods, specifically the ESM-1v model^24^ holds considerable promise for the identification of highly variable LRR domains. Building on ESM-LRR, we propose a framework named Plant Disease-Resistance Gene Predictor (R-Predictor) for the comprehensive annotation of plant R genes. This framework integrates our developed ESM-LRR method with leading methods for the identification of signal peptides (SignalP 6.0)^30^, transmembrane domains (TMHMM-2.0)^31^ and coiled-coil domains (Paircoil2)^32^, providing a robust and holistic solution for the annotation of R genes in plants.

We deconstructed the proteins coded by 445 experimentally verified R genes and obtained 15 distinct protein domain topologies with extensive coverage of recently characterised resistance genes. ESM-LRR demonstrated superior performance on an independent test set for the identification of LRR domains. The R-Predictor successfully identified RLKs, RLPs and NLRs simultaneously across the 15 different domain topologies, outperforming existing methods in the annotation of R genes in *Arabidopsis thaliana*. We also used R-Predictor to identify R genes in several crops including rice, tomato, and grapevine. Furthermore, given the recently released AlphaFold3^33^ demonstrated capability to predict protein-protein interactions and simulate 3D protein structures, we leveraged AlphaFold3 as a downstream method to screen 1116 protein pairs between the NLRs identified exclusively by our R-Predictor and relevant plant-pathogen effectors. This approach led to the identification of 15 candidate NLR-effector complexes. Our advancements in the development of the R-Predictor, combined with the application of AlphaFold3^33^ to validate our findings, dramatically enhance our understanding of plant immune system. These developments are expected to stimulate further research aimed at improving disease resistance in crops, for instance, through the application of genome editing or genomic selection to breeding disease resistant crops.

## Results

### Overview of ESM-LRR for the prediction of LRR domains

The ESM-LRR method was developed to enable the comprehensive prediction of LRR domains in protein sequences, using the deep protein language model ESM-1v^24^ (**Fig. 1a**) for domain characterisation in conjunction with the advanced machine learning method for regression and prediction. In this study, a total of 128,855 amino acid sequence fragments derived from 1,585 LRR-containing proteins from the Swiss-Prot database were used (**Supplementary Table 1**). Of these, 13,905 fragments were known as true LRR unites and were assigned the highest scores, while the remaining fragments were progressively assigned lower LRR scores based on their spatial distance from the authentic LRR units (**Extended Data Fig. 1**). The scores for each fragment were assigned according to a normal distribution probability density function (μ=0, σ=0.2). Although the scores of non-LRR units were not zero, they displayed significant differences from those of the true-LRR units, with higher scores indicating closer proximity to the true LRR units. These fragments were divided into a training set (80%) and a testing set (20%) to evaluate model performance.

**Fig. 1.**
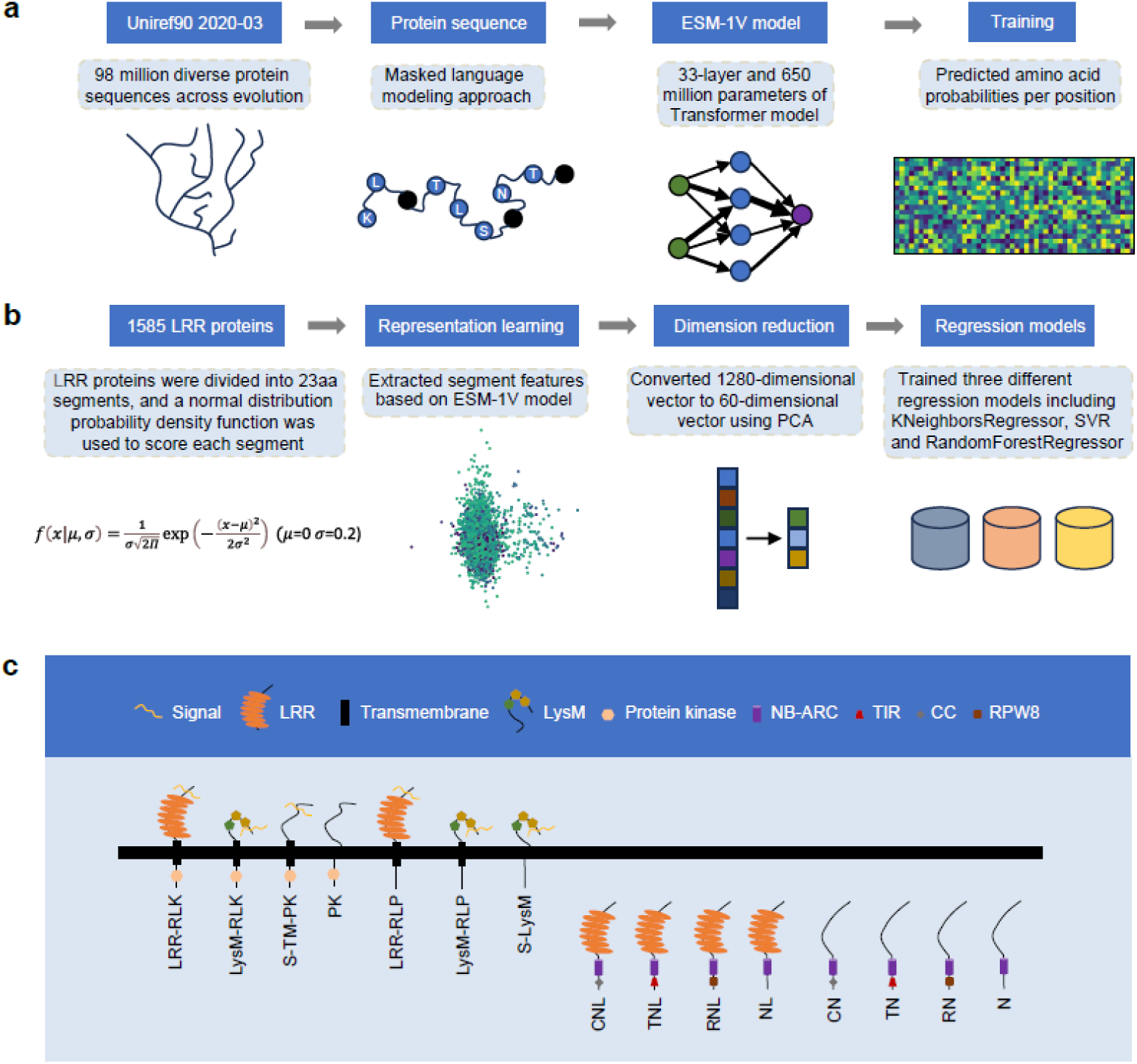
Overview of ESM-LRR and domain topologies of R genes. **a**, ESM-1v is a 650-million-parameter deep protein language model trained on 98 million protein sequences across all organisms. The model was trained using a masked language modeling task, where random residues are masked in the input sequences, requiring the model to predict the correct amino acid at each position, including the masked residues. **b**, ESM-LRR was trained on a dataset of 1,585 LRR proteins, using ESM-1v to generate sequence features, PCA dimensionality reduction, and inference combined with downstream machine learning models. **c**, Domain topologies of 445 experimentally verified disease-resistance gene products.

The LRR scores of the training set samples were decomposed into multiple dimensions (or features) using ESM-1v representation learning. These samples were subsequently visualized through Principal Component Analysis (PCA)^34, 35^ with each point colored according to its corresponding LRR score (**Fig. 2a**). The PCA plot visually indicated a separation trend, demonstrating that the differences between LRR and non-LRR units were captured by the features generated by ESM-1v. However, as PCA is an unsupervised method, it was unable to effectively discriminate between LRR and non LRR samples. Nonetheless, PCA efficiently reduced the high-dimensional matrix generated by ESM-1v representation learning into low dimensions, largely reducing computing time for downstream regression.

**Fig. 2.**
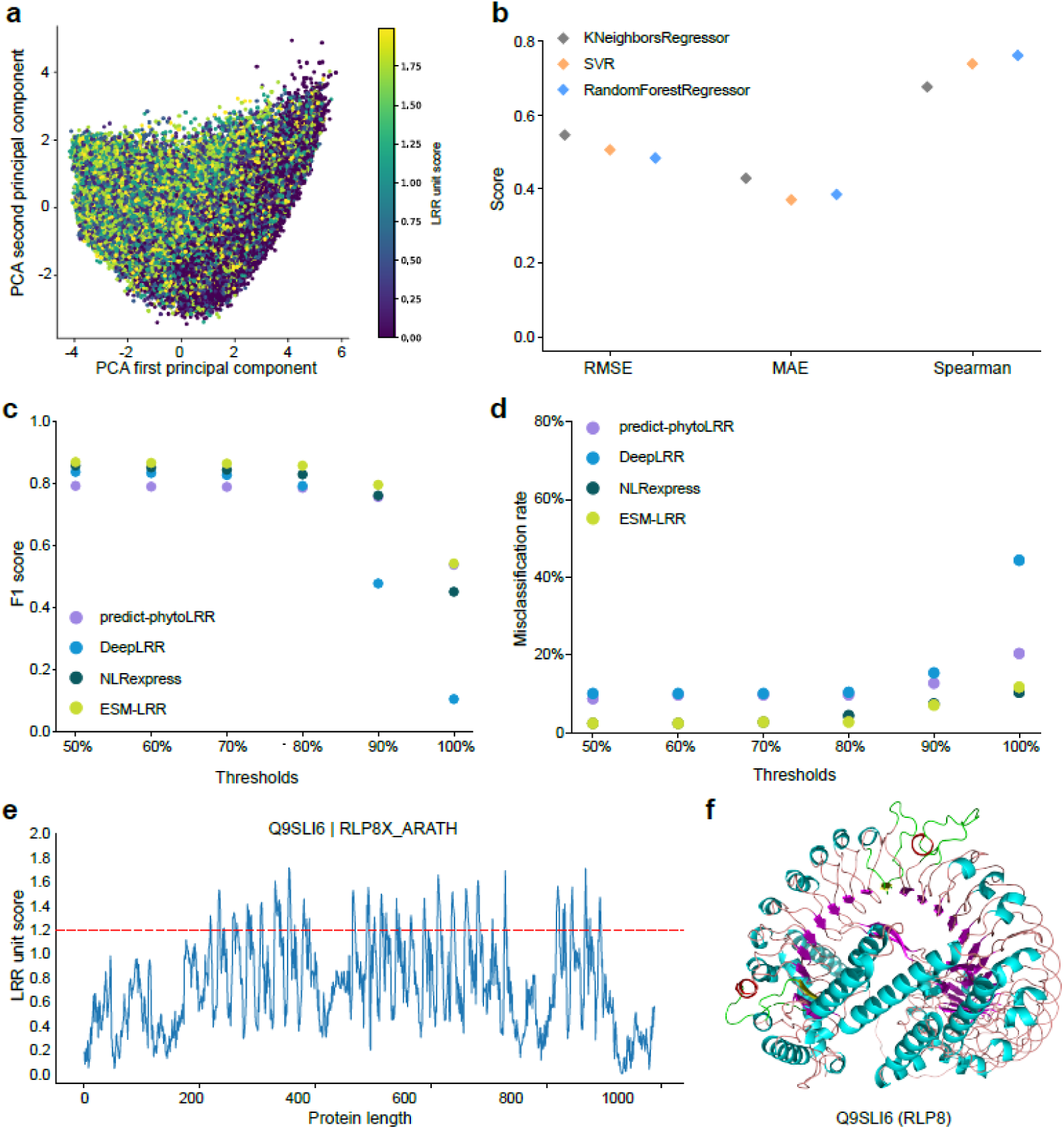
Comparison of ESM-LRR with other methods for annotating LRR domains. **a**, Training set was visualized after ESM-1v representation learning and PCA dimensionality reduction. **b**, The downstream machine learning models of ESM-LRR were tested based on an independent test set. **c-d**, Evaluation of the performance of ESM-LRR and other methods on 300 LRR proteins based on F1 score and misclassification rate. **e**, Q9SLI6 was divided using a sliding window size of 23 amino acids, and each fragment was assigned a score based on ESM-LRR. **f**, The 3D structure of Q9SLI6, with the LRR units that were not successfully annotated by ESM-LRR highlighted in red-green-yellow.

Three widely used machine learning methods - KNeighborsRegressor^36^, SVR^37^ and RandomForestRegressor^38^ were benchmarked, with only the most efficient method to be incorporated into the final version of ESM-LRR. (**Fig. 1b**). First, optimal hyperparameters were determined for each method based on five-fold cross-validation on the top 60 PCs of the representation matrix from ESM-1v, which mostly summarised the variance of the training dataset (**Supplementary Table 2**). These machine learning methods, each optimised with their respective hyperparameters, were then benchmarked using the test set.

RandomForestRegressor achieved scores of 0.48, 0.39 and 0.76 for Root Mean Squard Error (RMSE), Mean Absolute Error and Spearman Correlation Coefficient (Spearman), respectively. In comparison, KNeighborsRegressor produced scores of 0.55, 0.43 and 0.68, while SVR obtained scores of 0.51, 0.37 and 0.74 (**Fig. 2b** and **Supplementary Table 3**). No single model performed best across all metrics (the lowest RMSE, MAE, and the highest Spearman correlation). However, RandomForestRegressor was deemed the most optimal as it ranked first in two out of the three evaluation metrics (RMSE and Spearman). As a result, the final ESM-LRR method was constructed using ESM-1v representation learning combined with RandomForestRegressor.

### ESM-LRR outperforms existing methods in annotating LRR domains

To evaluate the performance of ESM-LRR in comparison to other methods, including predict-phytoLRR^39^, DeepLRR^20^ and NLRexpress^21^ in identifying LRR domains, we selected an additional set of 300 LRR proteins (comprising 2,812 LRR units) from the Swiss-Prot database (**Supplementary Table 1**). Six different matching thresholds (50%, 60%, 70%, 80%, 90% 100% identity) were established to define successful identification as true positives. F1 score and misclassification rate were used as metrics to evaluate the performance of these methods. ESM-LRR consistently achieved the highest F1 scores across all thresholds, with improvements ranging from 0.9% to 4.5% over the second-best method (**Fig. 2c** and **Supplementary Table 4**). When the matching threshold is 90% identity, ESM-LRR has the maximum improved F1 score of 0.80, while at the 100% threshold, it showed the smallest improvement with an F1 score of 0.54. The misclassification rate in this study was defined to determine the proportion of proteins containing LRR domains that were not correctly identified by the methods. Compared to other methods, ESM-LRR achieved the lowest error rate in five out of the six thresholds, except for the 100% threshold. At this threshold, NLRexpress missed 31 proteins with LRR domains, while ESM-LRR missed 35, corresponding to misclassification rates of 10.3% and 11.7 %, respectively (**Fig. 2d**).

One of the 300 LRR proteins, Q9SLI6 (putative receptor-like protein 8, RLP8), was used as a case protein to demonstrate the predictive capability of ESM-LRR. The assigned scores for each protein fragment are shown in **Fig. 2e**. Given that the flanking sequences of true LRR units often display considerable scores and present false positive peaks on the score curve, we developed a filtering script based on a fixed score threshold (>1.2) and the distance relationship between true LRR units, which can be easily specified by users (**Extended Data Fig. 2**). Q9SLI6 contains a large LRR domain, consisting of 26 LRR units. Among them, 23 were correctly identified by ESM-LRR (**Fig. 2f**). Although predict-phytoLRR and NLRexpress also identified 23 out of the 26 LRR units, they generated 1 or 3 false positive LRR units, respectively. Additionally, DeepLRR was able to identify only 3 LRR units. The development of ESM-LRR greatly improves the accuracy of LRR domain predictions in R genes in plants.

### The protein domain topologies of plant R genes

To gain a deeper understanding of domain topologies in plant R genes, we collected 445 experimentally validated genes encompassing various domain topologies^19, 40^ (**Supplementary Table 5**). Among these genes, approximately 6.7% (30/445) encode RLKs/RLPs, while approximately 93.3% (415/445) encode NLRs. RLKs and RLPs are membrane-bound and involved in PTI, whereas NLRs are intracellular proteins associated with ETI. Distinct domain topologies were observed between RLKs/RLPs and NLRs (**Fig. 1c**). Specifically, four domain topologies were identified in RLKs: (I) signal peptide-LRR domain-transmembrane domain-protein kinase domain (LRR-RLK); (II) signal peptide-LysM domain-transmembrane domain-protein kinase domain (LysM-RLK); (III) signal peptide-transmembrane domain-protein kinase domain (S-TM-PK); and (IV) protein kinase domain only (PK), (**Extended Data Fig. 3a-b**). The primary distinction between RLPs and RLKs is the absence of a protein kinase domain in the cytoplasmic region of RLPs. Three domain topologies were identified in RLPs: (I) signal peptide-LRR domain-transmembrane domain (LRR-RLP); (II) signal peptide-LysM domain-transmembrane domain (LysM-RLP); and (III) signal peptide-LysM domain (S-LysM) (**Extended Data Fig. 3a-b**).

In NLRs, six distinct domain topologies were identified: (I) NB-ARC domain-LRR domain (NL); (II) Toll/interleukin-1 receptor domain-NB-ARC domain-LRR domain (TNL); (III) coiled-coil domain-NB-ARC domain-LRR domain (CNL); (IV) Resistance to Powdery Mildew 8 (RPW8) domain-NB-ARC domain-LRR domain (RNL); (V) Toll/interleukin-1 receptor domain-NB-ARC domain; and (VI) RPW8 domain-NB-ARC domain (RN). Two additional domain topologies have also been reported in other annotation methods: (VII) NB-ARC domain (N); and (VIII) coiled-coil domain-NB-ARC domain (CN). However, these two were not present in the genes we collected, but have been included in our annotation framework (**Extended Data Fig. 3c-d**).

In summary, NLRs are characterized by the presence of NB-ARC domains, while RLKs are distinguished by the presence of protein kinase domains, both of which enable them to trigger downstream signalling, respectively. In contrast, RLPs lack such kinase domains and rely on RLKs or other proteins for signaling. Additionally, NLRs often contain TIR, RPW8 or coiled-coil domains at the N-terminus, which are absent in both RLKs and RLPs.

### Construction of the R-Predictor for the annotation of plant R genes

In addition to LRR domains, other critical domains, such as signal peptides, transmembrane domains, and coiled-coil domains, are also challenging to discern as highlighted in previous studies on domain topologies^30–32^. For instance, coiled-coil domains are particularly difficult to identify due to their composition of two or more α-helices arranged in parallel or antiparallel orientations. To achieve optimal identification of these complex domains, we selected commonly used methods for each domain and determined the most effective combination of these methods through comprehensive benchmarking. For the identification of signal peptides, LipoP^41^, Phobius^42^, Deepsig^43^ and SignalP 6.0^30^ were benchmarked using 9,796 proteins, each containing a single signal peptide from the Swiss-Prot database. SignalP 6.0 demonstrated the highest F1 scores and the lowest misclassification rates across all thresholds (**Figs. 3a, d** and **Supplementary Table 6**). For transmembrane domains, SCAMPI2^44^, HMMTOP^45^, Phobius and TMHMM-2.0^42^ were selected and benchmarked using 79,862 proteins containing a total of 381,188 transmembrane domains from the Swiss-Prot database. TMHMM-2.0 achieved the highest F1 scores across all matching thresholds (**Fig. 3b** and **Supplementary Table 7**). At thresholds of 50%, 60%, 70% and 80% identity, HMMTOP exhibited the lowest misclassification rates. However, at the 90% and 100% identity, TMHMM-2.0 yielded the lowest misclassification rates (**Fig. 3e, Supplementary Table 7**). For coiled-coil domains, we benchmarked Coils^46^, Paircoil2^32^, DeepCoil^47^ and CoCoNat^48^ using 3,783 proteins containing 6,438 coiled-coil domains from the Swiss-Prot database. Paircoil2 achieved the highest F1 scores and lowest misclassification rates across all maching thresholds (**Figs. 3c, f** and **Supplementary Table 8**).

**Fig. 3.**
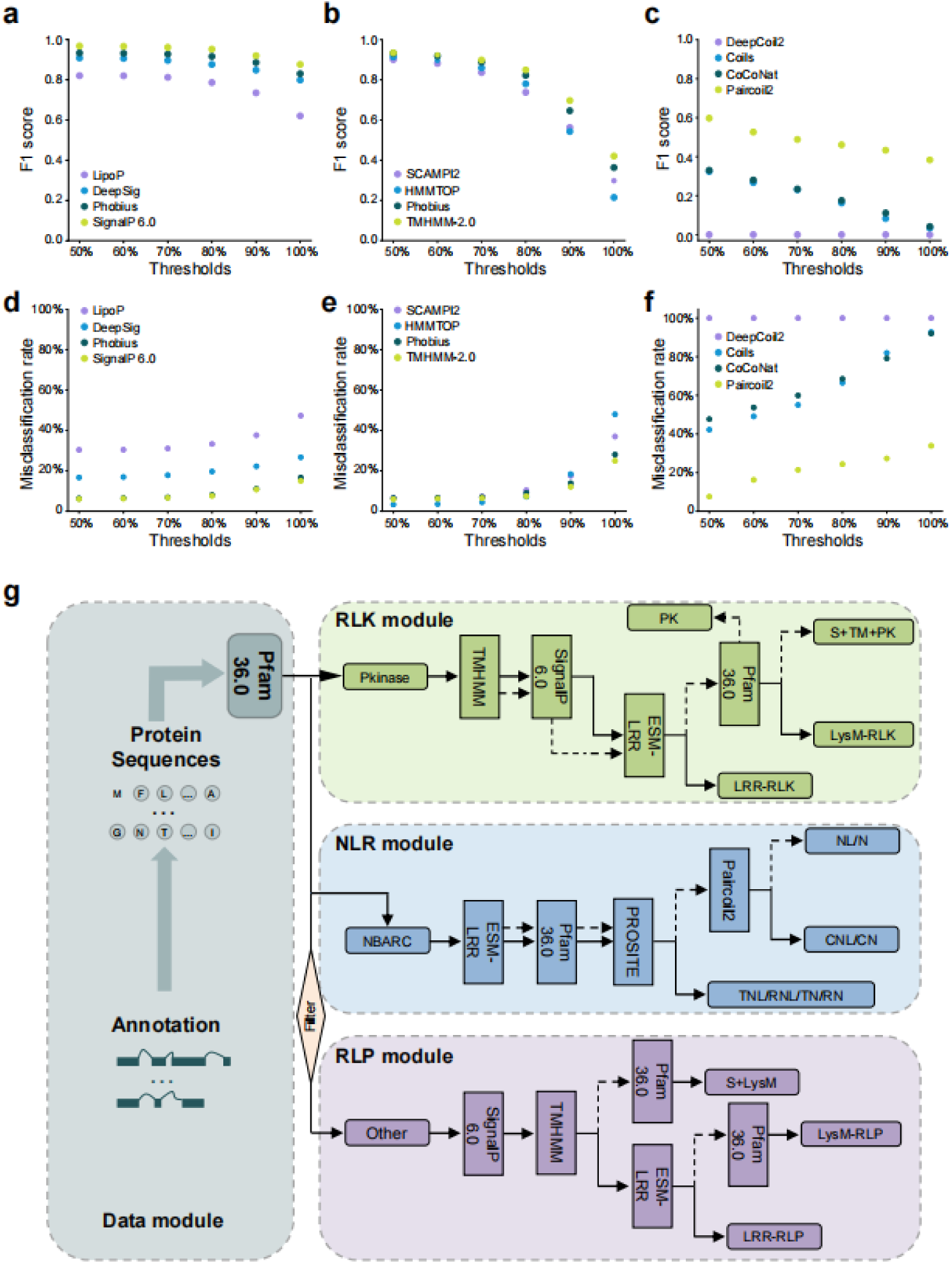
Benchmark various methods for different domains and overview of R-Predictor. **a-f**, the performance of different methods in annotating signal peptides, transmembrane domains, and coiled-coil domains was evaluated using F1 score and misclassification rate. **g**, R-Predictor consists of four modules. In the data module, protein sequences are classified and transmitted to three downstream modules. RLK module, NLR module and RLP module are responsible for outputting 4, 8 and 3 types of plant R genes, respectively.

The final framework, R-Predictor, designed for the de novo annotation of various R genes integrate four modules for data pre-processing and the identification of different types of proteins. Each module incorporates customized filtering scripts and the best-performing methods identified through benchmarking (**Fig. 3g**). Specifically, the Data module includes Pfam 36.0^49^, which can be flexibly integrated and can accurately determine whether a protein contains protein kinase domains, NB-ARC domains or neither. These two domains are highly conservative and robust in terms of identification methods. This module decides the appropriate subsequent module within the R-Predictor for identifying the input protein. SignalP 6.0 and TMHMM-2.0 are used in the RLK and RLP modules for the identification of signal peptides and transmembrane domains, while ESM-LRR is responsible for identifying LRR domains across all modules. In the NLR module, Pfam 36.0 and PROSITE^50^ work in conjunction to identify TIR and RPW8 domains, while Paircoil2 is used to detect coiled-coil domains. Overall, R-Predictor provides detailed information on the domain topologies of R genes along with the chromosomal locations of these genes when a corresponding genome annotation file is provided.

### Evaluation of R-Predictor across multiple crop species

To assess the performance of R-Predictor in comparison to existing tools, we annotated RLKs, RLPs and NLRs across functionally annotated genomes of Arabidopsis (Araport11)^51^, rice (IRGSP-1.0)^52^, tomato (ITAG4.0)^53^ and grape (PN40024_T2T)^54^. In contrast to RLKdb^22^, which exclusively identifies RLKs, annotated 285, 180, 186 and 139 RLKs for each species, R-Predictor identified a much higher number of RLKs with 1108, 791, 628 and 711 annotations respectively, while also detected 79, 92, 63 and 136 RLPs (**Fig. 4a**). For NLRs, NLRexpress^21^ annotated 273, 245, 113 and 470 NLRs, NLRtracker^19^ identified 420, 349, 212 and 581, Resistify^23^ annotated 421, 373, 212 and 746, while R-Predictor identified 422, 369, 212 and 745 NLRs in Arabidopsis, rice, tomato, and grape, respectively (**Fig. 4b**).

**Fig. 4.**
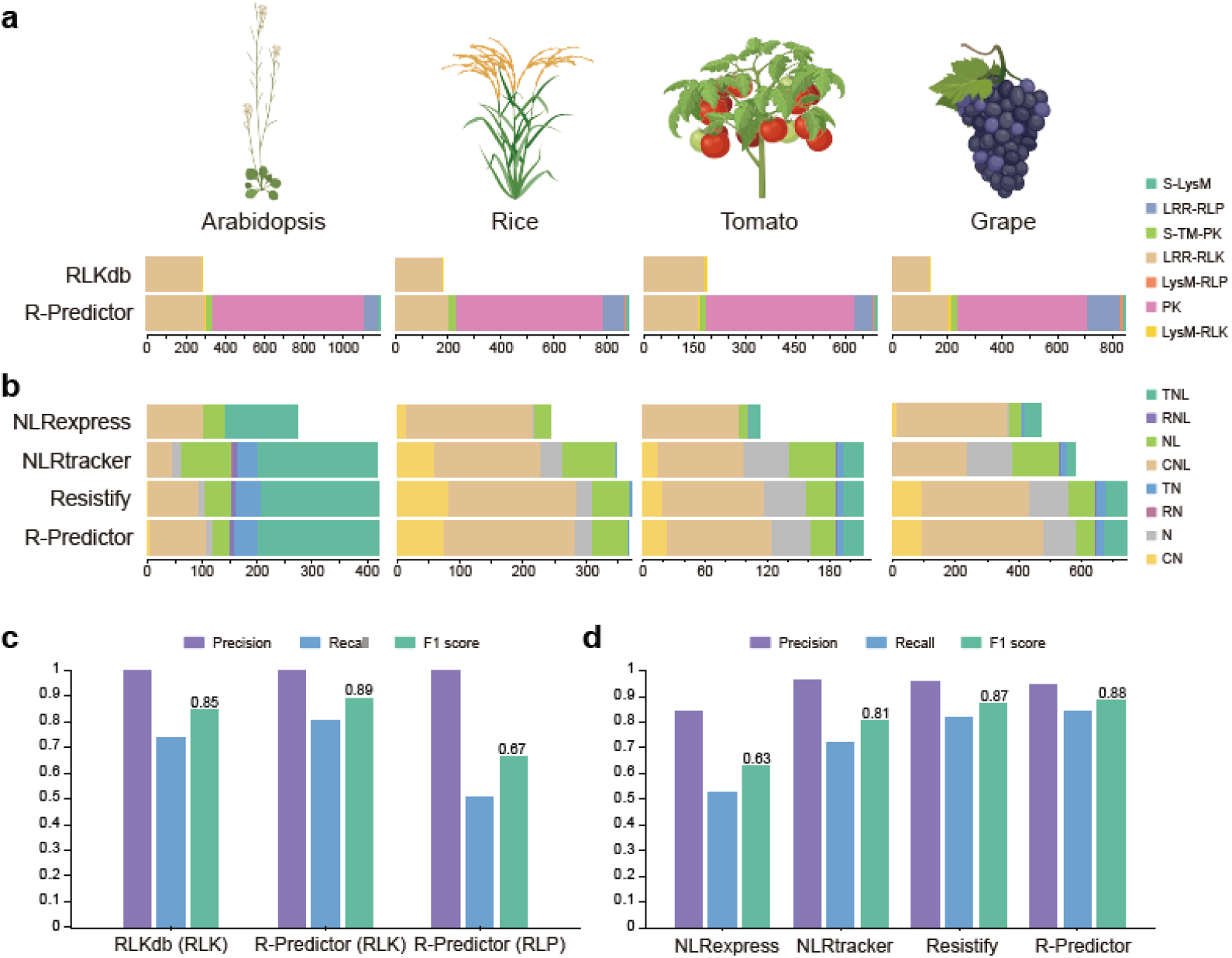
Comparison of R-Predictor with other methods for annotating R genes in various plant species. **a**, RLKs and RLPs with different domain topologies annotated by R-Predictor and RLKdb. **b**, NLRs with different domain topologies annotated by R-Predictor and other methods. **c**, Evaluation of the performance of R-Predictor and RLKdb to annotate RLKs and RLPs from Araprot11 based on precision, recall, and F1 score. **d,** Evaluation of the performance of R-Predictor and other methods to annotate NLRs from Araprot11 based on precision, recall, and F1 score.

Since the Araport11 contains high-quality and manually curated functional annotations, we further investigated its disease resistance gene annotations by comparing with the results from our framework and comparable methods. RLKdb identified only two types of RLK proteins: LRR-RLKs and LysM-RLKs. Both RLKdb and the R-Predictor identified 243 identical LRR-RLKs and 5 identical LysM-RLKs (**Extended Data Fig. 4a** and **Supplementary Tables 9-10**). Additionally, RLKdb identified 37 unique LRR-RLKs, of which 18 lacked the signal peptide, one lacked the protein kinase domain, two lacked the LRR domain and six contained only one or two LRR units. These six are highly likely to be false positives, as one or two LRR units are generally unstable. The R-Predictor achieved an F1 score of 0.89 for RLK annotation, representing a 4.8% improvement over RLKdb, largely due to the identification of 26 additional true LRR-RLKs (**Fig. 4c** and **Supplementary Table 11**). However, for RLPs, the R-Predictor obtained a less favorable F1 score of 0.67 (**Fig. 4c**). Upon examining the domain topologies of false negative proteins, we found that 53 LRR-RLPs annotated through general tools and manual curation lacked basic domain topologies: 44 lacked the signal peptide and nine lacked the transmembrane domain. Through R-Predictor, we identified 5 CNs, 103 CNLs, 11 Ns, 31 NLs, 2 RNs, 7 RNLs, 43 TNs and 220 TNLs in Arabidopsis thaliana, followed by Resistify which identified 4 CNs, 89 CNLs, 12 Ns, 48 NLs, 2 RNs, 7 RNLs, 45 TNs and 214 TNLs. Moreover, 0 CN, 46 CNLs, 16 Ns, 93NLs, 2 RNs, 7 RNLs, 38 TNs and 218 TNLs were identified by NLRtracker and 1 CN, 101 CNLs, 0 N, 40 NLs, 0 RN, 0 RNL, 1 TN and 130 TNLs were identified by NLRexpress (**Supplementary Table 12-15**). By generating the intersection of these results, 422 NLRs were identified by R-Predictor completely cover Resistify and NLRexpress, and there are 416 identical NLRs with NLRtracker. The confusion matrix shows that R-Predictor achieved the highest recall (0.84) and F1 score (0.88) across all methods, outperforming the second-place Resistify by 2.4% in recall and 1% in F1 score (**Fig. 4d** and **Supplementary Table 16**). The improvement can be attributed to the R-Predictor’s identification of 10 additional true NLRs, with the types of CNL (R-Predictor: 103 vs. Resistify: 89) and TNL (220 vs. 214) accounting for approximately half of the difference.

We also leveraged the evolutionary relationships of NLRs to investigate 46 NLRs, out of the 422 identified by the R-Predictor, that were not recorded as valid in Araport11 (**Extended Data Fig. 4b**). By constructing a phylogenetic tree, we found that the majority of NLRs (413/422) were successfully classified into evolutionary branches consistent with their identified types, except for 9 CNLs that appeared in the TNL branch. The misclassification is likely due to the high similarity between these two topologies, supporting the reliability of the 422 NLRs identified by R-Predictor. The 46 NLRs exclusively detected by R-Predictor (labeled in **Extended Data Fig. 4b)** were either embedded within or formed isolated branches adjacent to their corresponding identified branches. This indicates these NLRs are potential candidates for further study. However, experimental validation is required to confirm their functional roles. Similarly, NLR phylogenetic trees were constructed for rice, tomato and grapevine based on the 369, 212 and 745 NLRs annotated by R-Predictor, respectively (**Extended Data Figs. 5-7** and **Supplementary Tables 17-19**). The results were consistent with those obtained for Arabidopsis, further validating our method’s applicability across different plant species.

### AlphaFold3 screening identified 15 candidate NLR-effector complexes

To further evaluate the reliability of the 46 NLRs identified by R-Predictor but not annotated in Araport11, we employed AlphaFold3^33^ as a supplementary approach to determine whether any of these NLRs could form compatible complexes with experimentally verified plant-pathogen effectors. This analysis was based on the fact that NLRs typically interact with pathogen effectors to mediate effector-triggered immunity (ETI)^55–57^. We selected three of the shortest proteins from the 46 NLRs, including NP_001319997.1 (TNL/AT4G19500), NP_199537.2 (CNL/AT5G47260) and NP_001318809.1 (TN/AT5G56220), to reduce computing demand as limited by the AlphaFold3 web services. Additionally, we selected 372 known plant-pathogen effectors (PPEs) from PHI-base ^58, 59^, also taking protein length into consideration. These effectors are secreted by various plant pathogens, including *Pseudomonas syringae*, *Hyaloperonospora arabidopsidis*, *Ralstonia solanacearum*, *Magnaporthe oryzae*, *Phytophthora sojae* and 50 other plant pathogens.

A total of 1,116 protein pairs were tested for interactions between the three potential NLRs and the 372 plant-pathogen effectors. This analysis resulted in 15 pairs with a combined ipTM+pTM score of ≥ 1.2 (**Fig. 5a** and **Supplementary Table 20**). The ipTM+pTM score provided by AlphaFold3, ranges from 0 to 2 and serves as a confidence metric for both individual protein structures and their interactions within complexes, with higher values indicating greater confidence in prediction accuracy. Among the three NLRs, NP_001319997.1 was predicted to form a complex with G3C9P0 (ipTM+pTM = 1.22) (**Fig. 5b**), and this pathogen effector has been implicated in downy mildew from the oomycete pathogen *Hyaloperonospora arabidopsidis*^60^. NP_199537.2 was predicted to interact with A5CLR7 (ipTM+pTM = 1.29), A0A1I7RYM0 (ipTM+pTM = 1.24) and H9DUR1 (ipTM+pTM = 1.21), respectively (**Fig. 5c** and **Extended Data Fig. 8a-b**). These pathogen effectors come from *Clavibacter michiganensis*, *Bursaphelenchus xylophilus* and *Verticillium dahlia*. Previous studies have reported that A5CLR7 causes bacterial canker in tobacco^61^, A0A1I7RYM0 induces bacterial wilt in black pine^62^ and H9DUR1 is recognised by the cell-surface receptor Vel to prevent from vascular wilt diseases^63^. NP_001318809.1 was predicted to interact with 11 pathogen effectors, including A0A7W7P8L9 (ipTM+pTM = 1.41), Q8P4H6 (1.4), A0A7G5WGW9 (1.4), L8WVP3 (1.37), A0A2R2Z575 (1.36), N1J7E2 (1.35), D0P3S7 (1.25), A0A0A0S3X0 (1.25), G9J651 (1.21), G2X4U8 (1.21) and D0P1A8 (1.21) (**Fig. 5d** and **Extended Data Fig. 8c-l**). These 11 effectors originate from nine different plant pathogens, including *Xanthomonas campestris*, *Fusarium oxysporum*, *Rhizoctonia solani*, *Phytophthora capsici*, *Blumeria graminis*, *Phytophthora infestans*, *Leptosphaeria maculans*, *Pantoea stewartii* and *Verticillium dahlia*, and are associated with diseases such as bacterial leaf spot, black rot, sheath blight, foliar blight, canker, anthracnose, powdery mildew, late blight, and bacterial wilt diseases^64–73^. While experimental validations of these 15 predicted complexes are required, AlphaFold3 demonstrates itself as an effective tool that complements R-Predictor, offering insights into the mechanisms underlying plant immune responses.

**Fig. 5.**
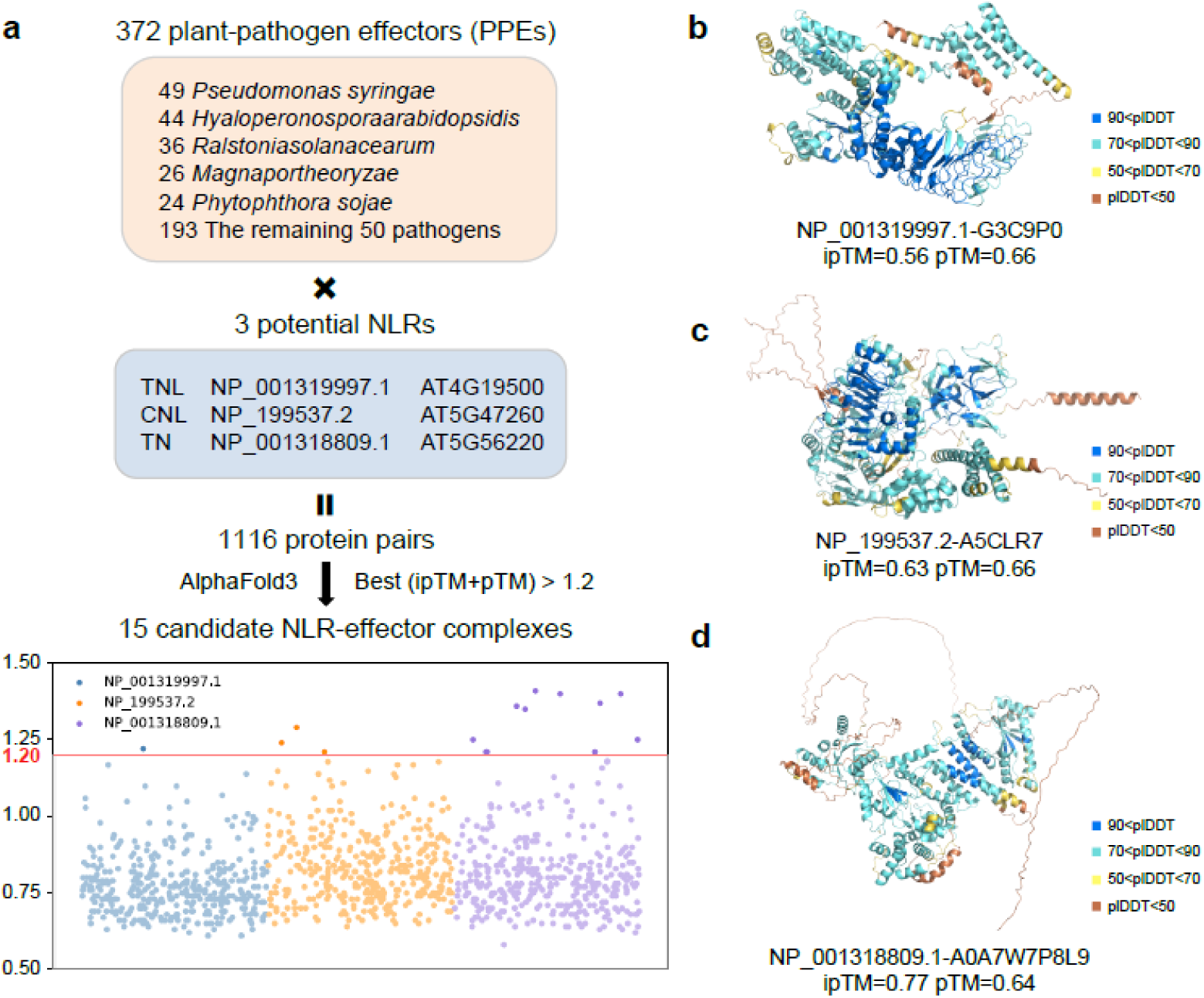
AlphaFold3 screen between 3 NLRs and 372 effectors identifies 15 candidate NLR-effector complex. **a**, Using AlphaFold3, 372 effectors from 55 plant pathogens were screened to identify complexes with three NLRs annotated by R-Predictor. Protein pairs with best ipTM+pTM score of 1.2 or higher were selected from the 1,116 protein pairs, resulting in 15 putative complexes. The best ipTM+pTM scores for all 1,116 complexes listed on the bottom. **b-d**, Three-dimensional structures of the complexes with the highest best ipTM+pTM scores by screening the complexes of NP_001319997.1, NP_199537.2 and NP_001318809.1 with 372 effectors. plDDT is used to determine the confidence of each residue in the predicted protein structure, with values ranging from 0 to 100.

## Discussion

Plants are constantly exposed to a wide range of pathogens, including viruses, bacteria, fungi, oomycetes and parasites, which can cause diseases when the immune defenses of plants are compromised^6^. Therefore, understanding the molecular mechanisms underlying plant immunity is essential for enhancing their defense against these threats. Unlike animals, which possess both innate and adaptive immune systems, plants rely exclusively on innate immunity to recognize and respond to pathogens^74–76^. Plant immunity is broadly classified into two types: PTI, which is primarily mediated by RLKs and RLPs, and ETI, which is mainly governed by NLRs. Recent advancements in sequencing technologies have facilitated the exploration of wild germplasms, revealing a wealth of disease-resistance traits beyond the scope of individual reference genomes^77–80^. Therefore, there is an urgent need for more precise and comprehensive annotation tools for identifying plant R genes, such as RLKs, RLPs and NLRs.

The LRR domain is a critical block present in the majority of RLKs, RLPs and NLRs, which influences accurate annotation^81^. Due to the diverse functions of LRR domains, involving in both plant immunity and a range of biological processes^82^, they exhibit high sequence variability. Current methods for annotating LRR domains include: predict-phytoLRR based on the position-specific scoring matrix (PSSM) algorithm^39^, DeepLRR based on convolutional neural networks (CNN)^20^, and NLRexpress based on a combination of machine learning predictors^21^. In contrast, our proposed ESM-LRR method leverages the deep protein language model ESM-1v^24^ to generate high dimensional features that provide a detailed characterisation of LRR units, while utilizing the machine learning model RandomForestRegressor^38^ to learn from these features and predict the probability that a new sequence contains LRR units. Compared to predict-phytoLRR^39^ and NLRexpress, which rely on pre-defined patterns such as “LxxLxLxxN” and “LxxLxL” for the recognition of LRR domains, both DeepLRR^20^ and ESM-LRR are more data driven by extracting features directly from known LRR units for training, and predicting LRR domains based on these trained models. Of the two methods, DeepLRR uses classification models to predict LRR units, whereas ESM-LRR employs regression models to estimate the probability of LRR units. These methodological differences may contribute to ESM-LRR’s superior performance compared to existing methods. However, there are areas in which ESM-LRR can be further improved. Due to computing limitations, the input and output of ESM-LRR are currently restricted to 23 amino acid residues in length. Although this is the most common length for LRR units (**Extended Data Fig. 9**), it imposes a constraint that falls short of broader expectations for more comprehensive analyses.

Compared to existing methods for annotating plant R genes^19–23^, our proposed R-Predictor has two major improvements: 1) it simultaneously annotates 15 distinct types of domain topologies for a wide range of R genes on a genome-wide scale. R-Predictor can also identify the functional genes lacking typical domain topologies, such as CERK1^13^, EDR1^14^, CEBiP^13, 15^, NRG1^16^ and DAR5^17^, providing an opportunity to recover previously overlooked resistance resources. 2) R-Predictor integrates the most efficient methods for identifying specific functional domains, such as SignalP 6.0^30^ for signal peptides, TMHMM-2.0^42^ for transmembrane domains, Paircoil2^32^ for coiled-coil domains, and ESM-LRR for LRR domains, thereby enhancing its ability to accurately classify R genes. For instance, R-Predictor demonstrates higher accuracy in identifying signal peptides, transmembrane domains and LRR domains compared to RLKdb^22^ and/or the general annotation in Araport11^51^. Additional, based on ESM-LRR and Paircoil2, R-Predictor can annotate more CNLs than other methods focused on NLRs identification. However, like most methods for annotating plant R genes, R-Predictor still relies on traditional tools such as Pfam 36.0^49^ and InterProScan^83^ to identify conserved protein kinase and NB-ARC domains, which may limit the discovery of novel R genes. Recent studies have demonstrated that deep learning approaches outperform traditional methods in identifying conserved protein domains^84, 85^, but the practical application of these advanced techniques still requires further refinement.

For validating the results of R-Predictor, AlphaFold3^33^ was employed to uncover previously unknown interactions between potential disease-resistance proteins and already reported pathogen-effectors. Since AlphaFold3 is currently available only as a web service, we have not yet implemented large-scale interaction prediction within our framework. Nevertheless, this approach advances our knowledge of the interaction dynamics between resistance genes and pathogen-effectors.

The most effective and sustainable strategy for mitigating crop diseases remains the incorporation of R genes into crop varieties. Our developed R-Predictor for resistance genes identification, coupled with the use of AlphaFold3 as a complementary method for the identification of pathogen effector-resistance protein interactions, represents a massive advancement in understanding resistance mechanisms. These efforts will also support future strategies for deploying R genes in breeding programs or through gene stacking for more robust disease resistance in crops.

## Methods

### Datasets

We screened 1,885 curated proteins containing LRR domains from the Swiss-Prot database (https://ftp.ncbi.nlm.nih.gov/blast/db/FASTA/) to train, verify and test ESM-LRR. 1,585 proteins were used for the development of ESM-LRR, and 300 proteins were applied for comparison of the performance of ESM-LRR and other methods for predicting LRR domains. We extracted 23aa sequence fragments along both sides of the LRR unit of 1,585 LRR proteins, and a normal distribution density function (=0, =0.2) was employed to assign scores for these fragments, ultimately generating a set consisting of 128,855 fragments, of which 13,905 fragments are true LRR units. Referring to the division ratio of 80%:20%, 103,084 fragments are responsible for 5-fold cross-validation to obtain the optimal hyperparameter combination of ESM-LRR, and 25,711 fragments are used as a test set. In addition to this, we also generated a signal peptide dataset, a transmembrane domain dataset and a coiled-coil dataset, which supported benchmarking of different modules in the R-Predictor. Proteins in these datasets are also from the Swiss-Prot database. The signal peptide dataset consists of 9,796 proteins, including 9,796 signal peptides, the transmembrane domain dataset consists of 79,862 proteins, including 381,188 transmembrane domains, and the coiled-coil dataset consists of 3,783 proteins, of which 6,438 coiled-coils.

### Training ESM-LRR

The training process of ESM-LRR was divided into the following steps, (I) sequence embeddings were extracted from 103,084 training fragments based on the deep protein language model ESM-1v^24^ (version: esm1v_t33_650M_UR90S_1); (II) loaded sequence embeddings and converted 1280-dimensional sequence embeddings to 60 dimensions by PCA^34, 35^; (III) 5-fold cross-validation and grid search to explore the optimal hyperparameter combination corresponding to KNeighborsRegressor^36^, SVR^37^ and RandomForestRegressor^38^; (IV) an independent test set was used to determine the final downstream model of ESM-LRR. The development of ESM-LRR was implemented on torch (version: 2.3.0) and scikit-learn (version: 1.5) frameworks, and GPU is GeForce RTX 3090 24GB.

### Identifying the LRR domain by ESM-LRR

If ESM-LRR only loads the downstream model, it can only assign scores to sequence fragments to recognize whether they are potential LRR units, but cannot identify the complete LRR domain. To improve the practicality of ESM-LRR, we incorporated a filtering script based on a fixed threshold and the distance relationship between LRR units. This script will filter the result given by the downstream model through three steps in sequence. (I) Remove sequence fragments with scores less than a given threshold, for example, threshold is 1.2 in this study. Regarding the fixed threshold set in the filtering script, we found in cross-validation that a threshold of 1.2 provides the best trade-off between precision and recall. (II) Pick a sequence fragment with a higher score if the distance between the two fragments is less than 20 amino acid residues. (III) If the distance between a fragment and its adjacent fragments is less than or equal to 30 amino acid residues, the fragment is considered as a potential LRR unit.

### Evaluation measures

We adopt 10 measures to evaluate the performance in this study, 4 for training ESM-LRR, 2 for benchmarking different modules of R-Predictor and 4 for prediction of R genes. To facilitate the description, hereafter, *y_i_* represents the actual value, *ŷ_i_* represents the predicted value, *y̅* represents the mean of actual values, *d_i_* represents the difference between the ranking of i-th pair of actual value and predicted value, and n represents total number of samples.

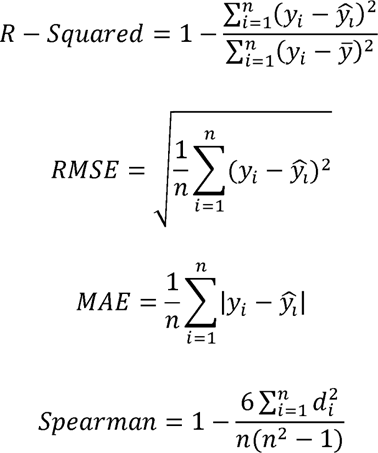

F1 score and misclassification rate are responsible for benchmarking different modules in R-Predictor. They are defined as follows, where *TP* means the number of actual values correctly identified, *FP* means the number of non-actual values incorrectly identified as actual values, *TN* means the number of non-actual values correctly identified and *FN* means the number of actual values incorrectly identified as non-actual values.

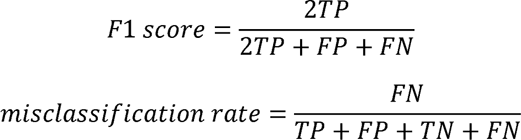

As predicting R genes is a multi-classification task, *Accuracy*, *Precision_Weighted_*, *Recall_Weighted_* and *F1 score_Weighted_* are used to evaluate the performance of various methods. They are defined as follows, where *t* means the number of different labels and *w_i_* 523 means the proportion of the number of samples corresponding to the i-th label to the total number of s

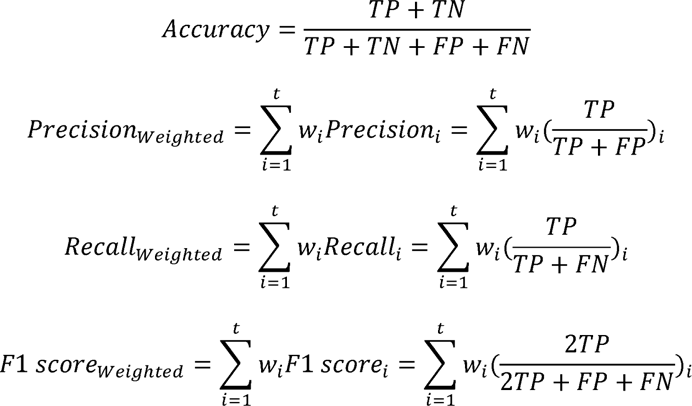

### Configuring the various programs for benchmarking

Predict-phytoLRR (https://github.com/phytolrr/predict-phytolrr), DeepLRR (version: 1.01 https://github.com/zhenyaliu77/DeepLRR) and NLRexpress (https://github.com/eliza-m/NLRexpress) were installed to identify LRR domain from 300 LRR proteins. LipoP (https://services.healthtech.dtu.dk/services/LipoP-1.0), DeepSig (https://deepsig.biocomp.unibo.it), Phobius (https://phobius.sbc.su.se) and SignalP 6.0 (https://github.com/fteufel/signalp-6.0) were implemented to participate in the benchmarking of signal peptide based on 9,796 protein sequences. SCAMPI2 (https://github.com/ElofssonLab/scampi2), HMMTOP (http://www.enzim.hu/hmmtop), Phobius (https://phobius.sbc.su.se) and TMHMM-2.0 (https://services.healthtech.dtu.dk/services/TMHMM-2.0) were used to identify the transmembrane domain from 79,862 proteins for benchmarking. In the benchmarking of coiled-coil, DeepCoil (version: 2.0 https://github.com/labstructbioinf/DeepCoil), COILS (version: 2.2 https://mybiosoftware.com/coils-2-2-prediction-coiled-coil-regions-proteins.html), CoCoNat (https://github.com/BolognaBiocomp/coconat) and Paircoil2 (https://cb.csail.mit.edu/cb/paircoil2) were employed to process 3,783 proteins. To present the performance of each program fairly, only default parameters are allowed in the benchmarking.

### Construction of the R-Predictor

From the benchmarking results, we determined a series of methods that presented high precision, recall and F1 score. Integrating these methods, we have developed a pipeline called R-Predictor, which combines the best-performing methods and subsequent filtering scripts for de novo identification of different types disease-resistance gene. R-Predictor is deployed on the Linux system in a modular way, each method has its own virtual environment to facilitate the user to update and maintain the pipeline. The operation of R-Predictor can be summarized as the following steps. (I) Loading a protein file and check whether it conforms to the fasta format. (II) Running pfam_scan.pl to search the Pfam 36.0 database^49^ to identify protein sequences that have either PK domain (PF00069, PF07714) or NB-ARC domain (PF00931) or neither. In addition, TMHMM-2.0^42^ was executed to identify protein sequences that contain both PK domain and TM domain. (III) SignalP 6.0^30^ was used to identify signal peptides in RLKs and RLPs, and TMHMM-2.0 was used to examine TM domains in RLPs. (IV) Implementing ESM-LRR to identify potential LRR-RLKs and LRR-RLPs, as well as to obtain protein sequences without the LRR domains for next step. (V) Running pfam_scan.pl again to identify LysM domain (PF01476), filtering out non-target domains and determining potential LysM-RLKs, LysM-RLPs and protein sequences that present the three domain topologies of S-TM-PK, PK and S-LysM. (VI) The Pfam 36.0 and PROSITE databases^49, 50^ were search to identify protein sequences containing TIR domain (PF01582, PF13676, PF18567, PS50104) or RPW8 domain (PF05659, PS51153) based on pfam_scan.pl and ps_scan.pl, respectively, and TNLs, TNs, RNLs and RNs were returned. Next, Paircoil2^32^ was applied to protein sequences that contain only NB-ARC domain or both NB-ARC domain and LRR domain to detect the presence of CC, and CNLs, CNs, NLs and Ns were returned. (VII) If the genome annotation file is provided, an embedded script will automatically return the gene ID and its chromosome position corresponding to the proteins.

### Execution of current methods for identifying R genes

RLKdb (https://biotec.njau.edu.cn/rlkdb) is a database containing ∼220,000 RLKs from 300 plant genomes, from which we screened LRR-RLKs belonging to Araport11^51^, IRGSP-1.0^52^, ITAG4.0^53^ and PN40024_T2T^54^. NLRexpress (https://github.com/eliza-m/NLRexpress), NLRtracker (version: v1.0.3 https://github.com/slt666666/NLRtracker) and Resistify (https://github.com/SwiftSeal/resistify) were deployed on the Linux system (Ubuntu 20.04.4) to quickly and large-scale identify NLRs in Araport11, IRGSP-1.0, ITAG4.0 and PN40024_T2T. In-house developed Python scripts were used to collate and evaluate the results produced by these methods.

### Phylogenetic tree

The sequence of NB-ARC domain in NLRs were delivery to MEGA^86^ (version: 11.0.13 https://www.megasoftware.net) for multiple sequence alignment and phylogenetic tree construction, and the default parameters were used. ITOL^87^ (version: v6 https://itol.embl.de) was employed to add annotations to the phylogenetic tree, including modifying color and labeling.

### NLR-effector complex prediction with AlphaFold3

NLR-effector complexes were modeled using AlphaFold3^33^. As the developer of AlphaFold3 did not provide model code to the academic community, we applied its website service (https://golgi.sandbox.google.com) to complete the prediction of NLR-effector complexes. It should be considered that due to website restrictions, each account can only submit 20 tasks per day, which is extremely unfriendly to the large-scale task. pTM is an integrated measure of AlphaFold3’s performance in predicting the overall structure of a complex, representing the predicted TM score for the superposition of the predicted structure with the hypothetical true structure. A pTM score above 0.5 indicates that the overall predicted fold of the complex is likely to resemble the true structure. ipTM measures the accuracy of predicted relative positions of subunits in a protein-protein complex, with values between 0.6 and 0.8 indicating a gray area where predictions could be either correct or incorrect. In practice, the overall confidence of multimer predictions should be based on the combination of pTM and ipTM, so we delineate multimers with a pTM plus ipTM greater than or equal to 1.2 as reliable. Finally, Pymol was used to draw the three-dimensional structure of proteins^88^.

## Data availability

The raw data in this study have been deposited into Github: https://github.com/zhouyflab/R-Predictor

## Code availability

All scripts performed in this study are available on Github: https://github.com/zhouyflab/R-Predictor

## Acknowledgements

We thank members of the Zhou lab at AGIS for discussion and comments on the project. This work was supported by the National Key Research and Development Program of China (No. 2023YFD2200700; 2023YFF1000100) and the Science Fund Program for Distinguished Young Scholars of the National Natural Science Foundation of China (Overseas) to Yongfeng Zhou.

## Author contributions

Y.Z. and Y.W. designed the project. Y.Z. and Y.W. supervised the project. Z.L. designed and trained the ESM-LRR. Z.L., X.W., S.C. and Z.L. performed the benchmarks. Z.L. designed and constructed the R-Predictor. Z.L. performed the bioinformatic analyses. Z.L., X.W. and T.L. prepared the figures and supplementary data. Z.L. wrote the draft. Y.Z. and Y.W. revised the manuscript. All authors contributed to manuscript preparation and read, commented on, and approved the manuscript.

## Competing interests

The authors declare no competing interests.

## Notes

### Competing Interest Statement

The authors have declared no competing interest.

