## Supplemental Figures for "Deep learning facilitates precise identification of disease-resistance genes in plants"

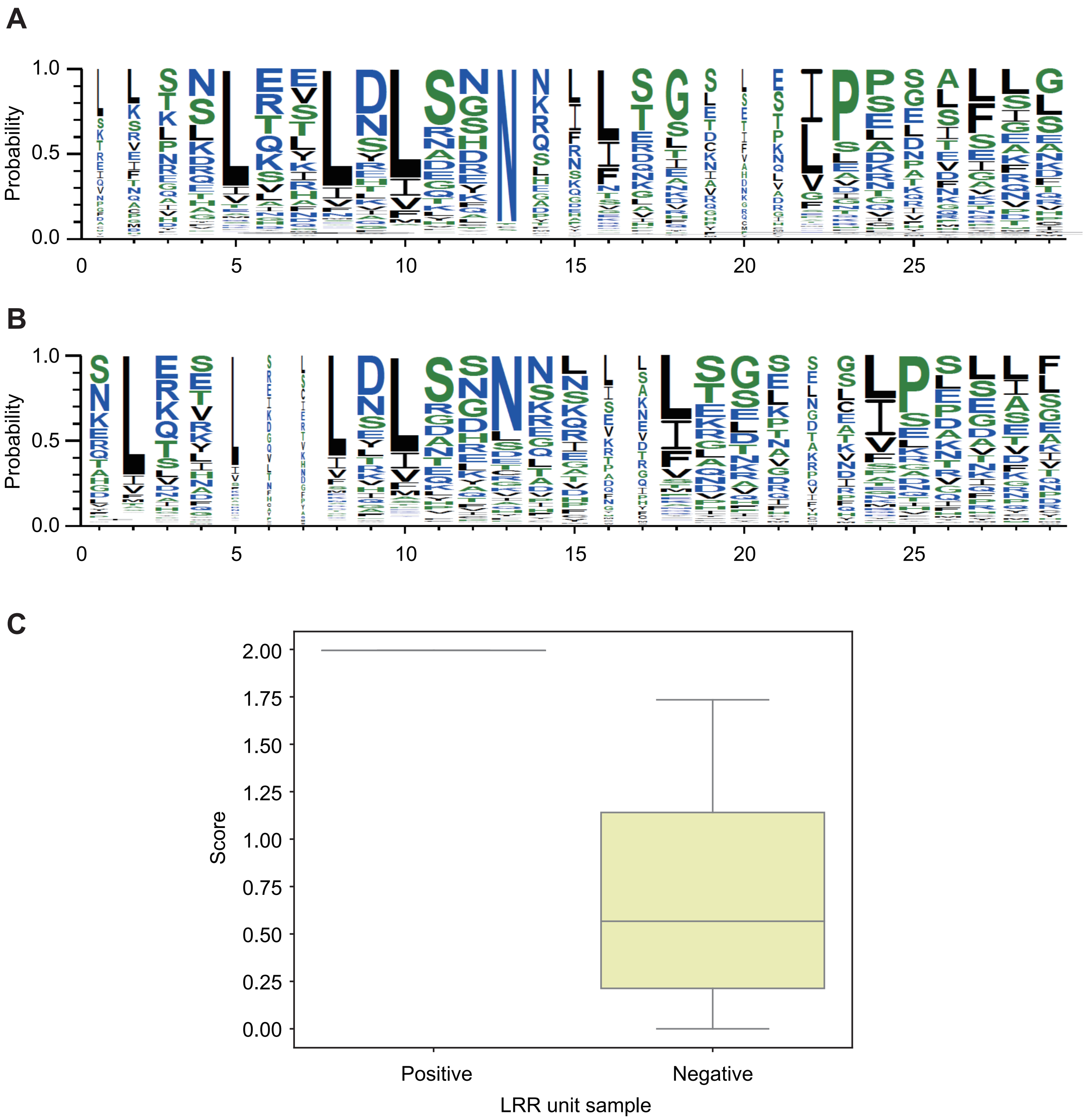


**Extended Data Fig. 1** | A. The conserved short sequence fragment pattern of 13,905 positive samples. B. The conserved short sequence fragment pattern of 13,905 samples randomly selected from 114,950 negative samples. C. Box plot of the scores of positive and negative samples in the ESM-LRR training set.

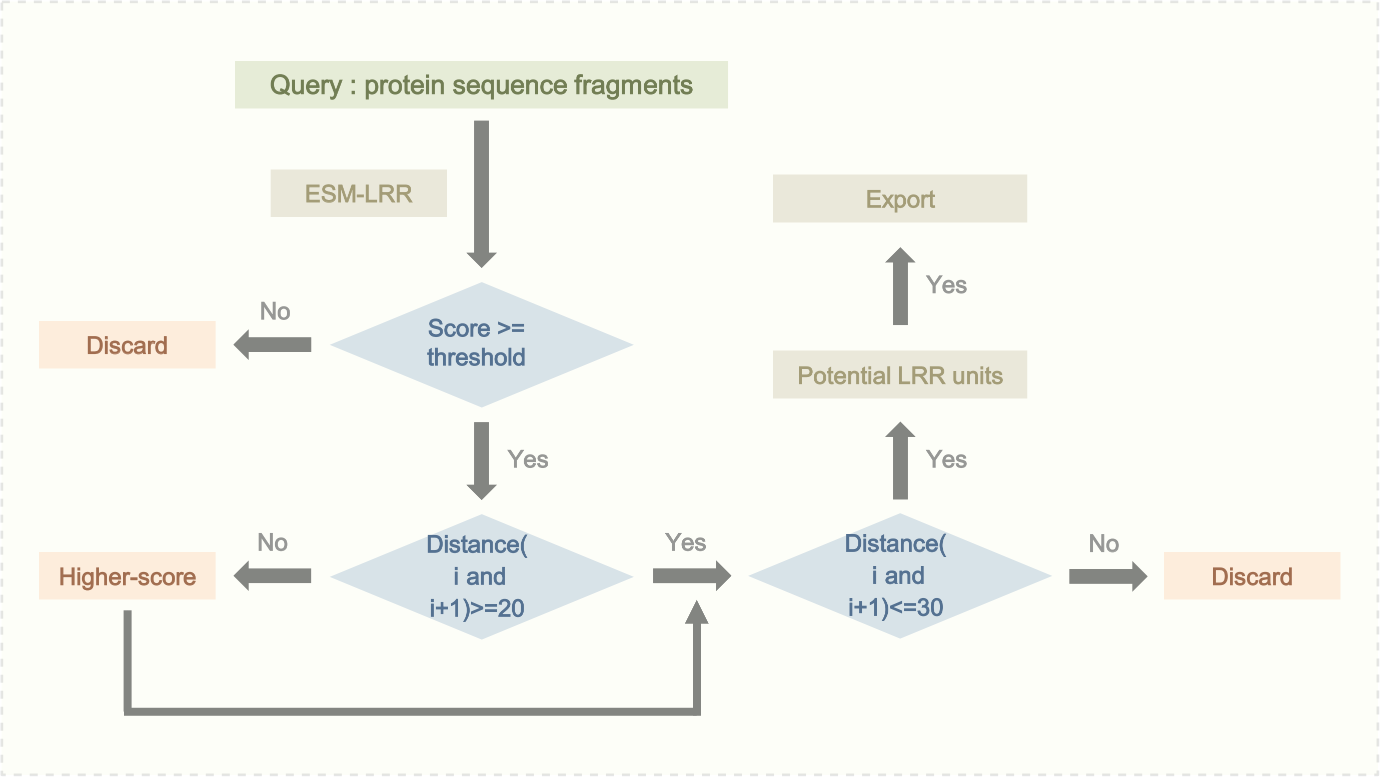


**Extended Data Fig. 2** | Overview of the filtering script that processes the output of ESM-LRR.

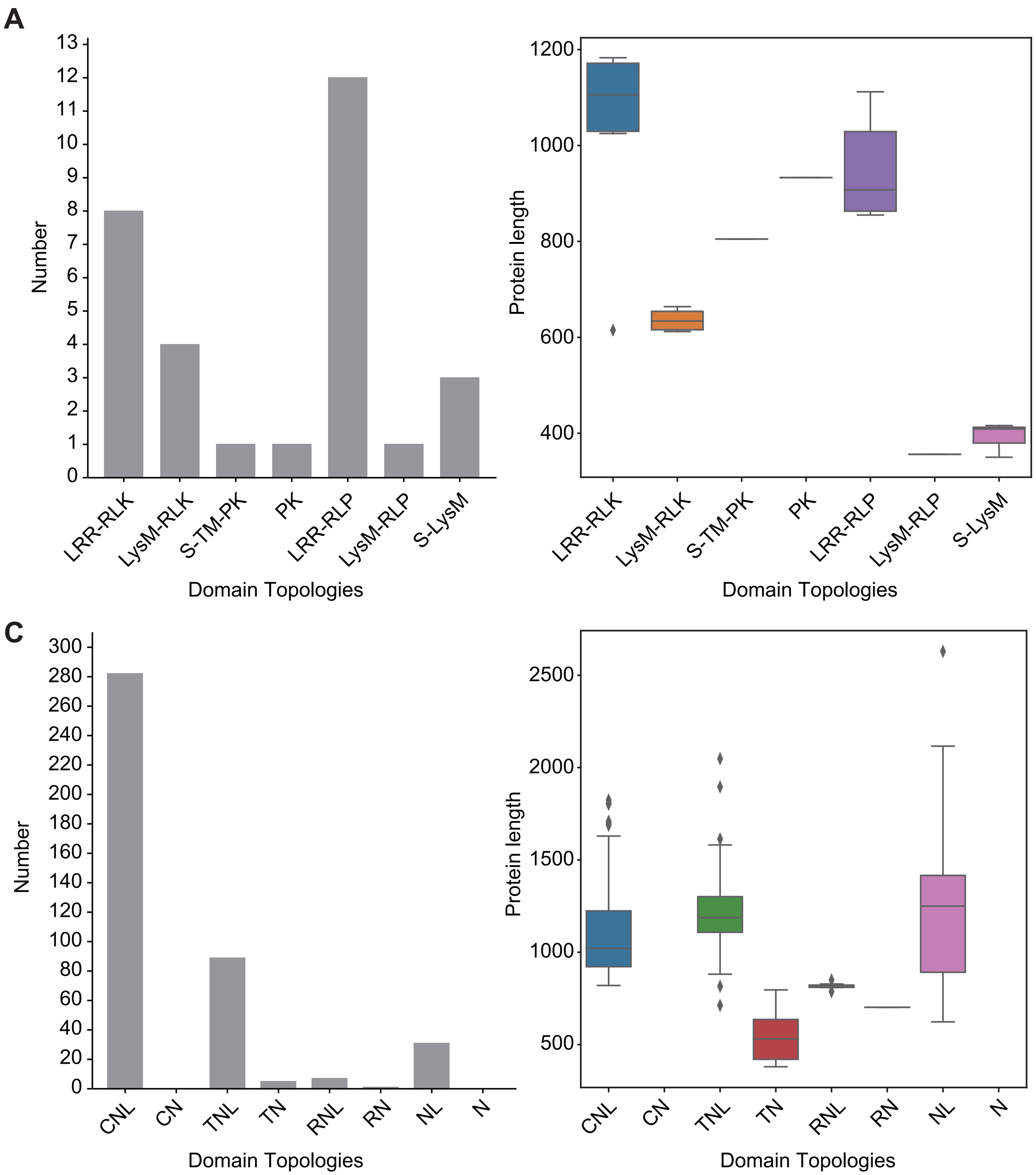


**Extended Data Fig. 3** | A. Number distribution of RLKs and RLPs in various domain topologies. B. Length distribution of RLK/RLP with different domain topologies. C. Number distribution of NLRs in various domain topologies. D. Length distribution of NLR with different domain topologies.

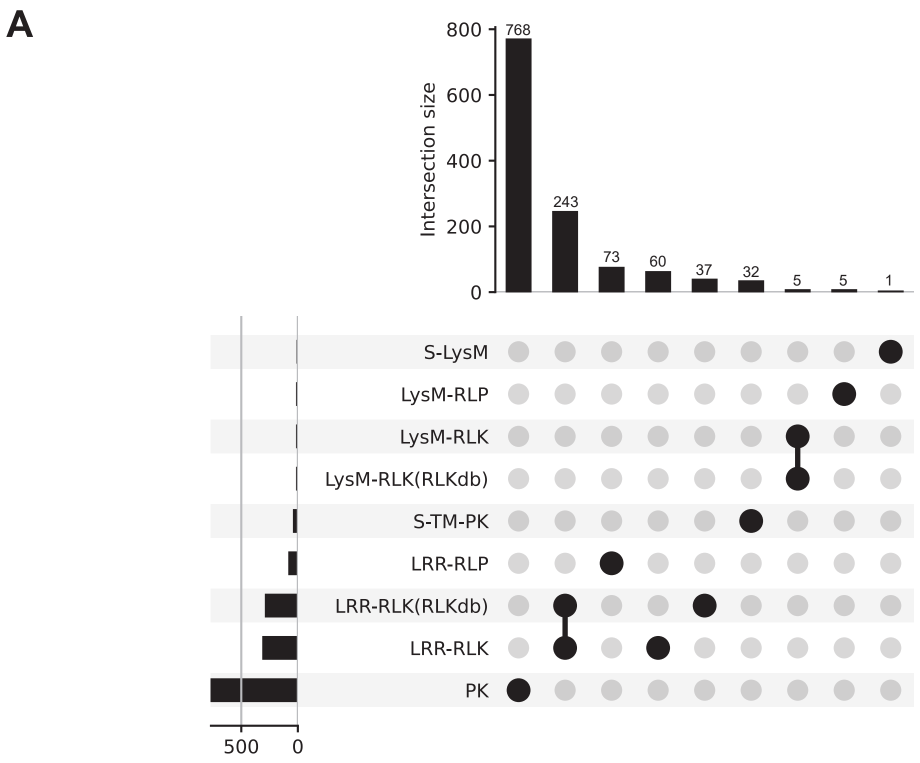


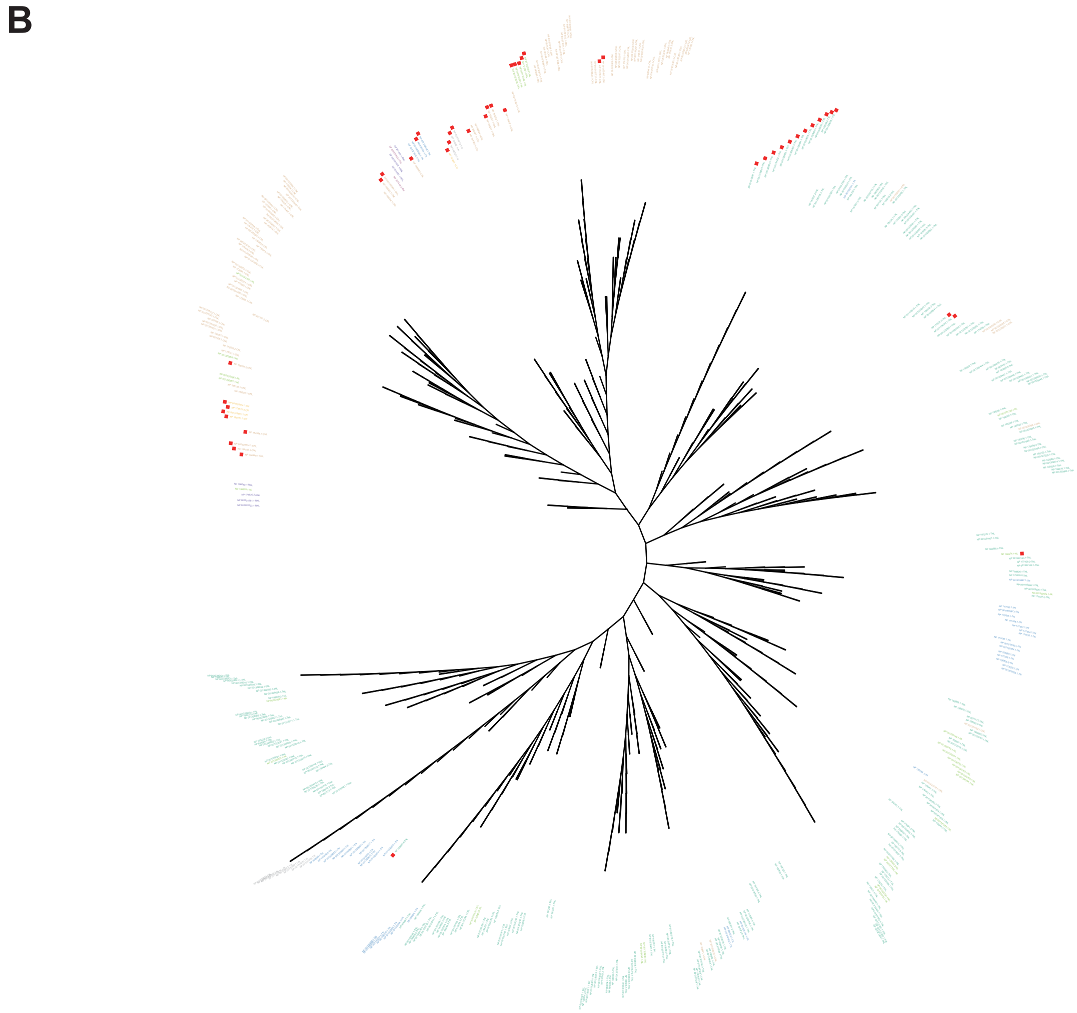


**Extended Data Fig. 4** | A. Upset plot of R-Predictor and RLKdb annotation results in Araport11. B. A phylogenetic tree constructed based on 422 NLRs annotated by R-Predictor. Leaf nodes are marked with different colors, representing NLRs with different domain topologies. NLRs marked by red rectangles cannot be annotated by Araport11.

**A**

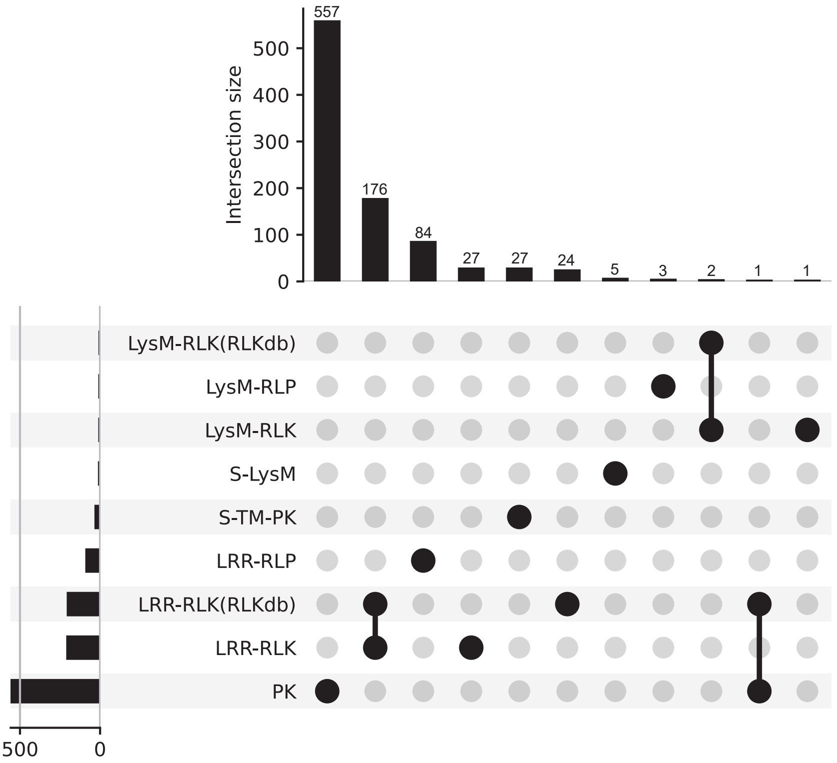


**B**

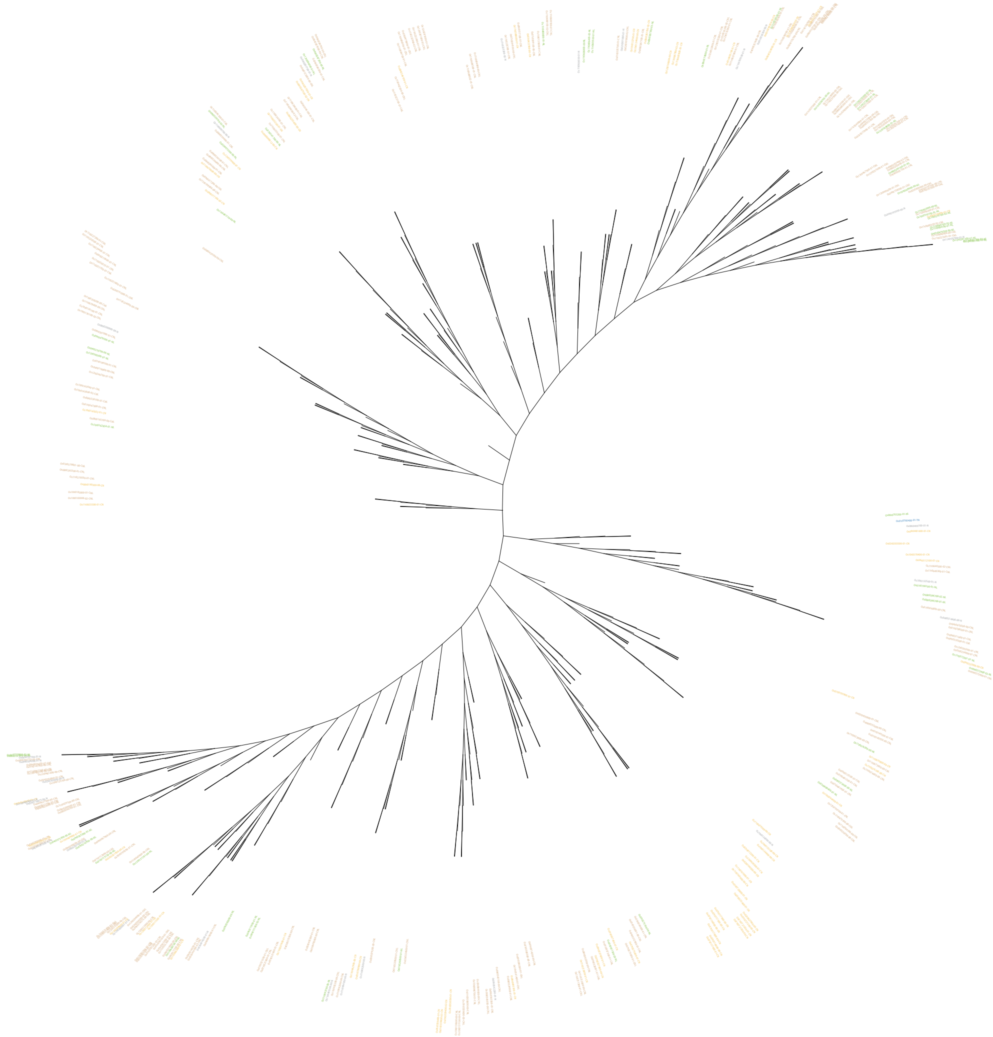

**Extended Data Fig. 5** | A. Upset plot of R-Predictor and RLKdb annotation results in IRGSP-1.0. B. A phylogenetic tree constructed based on 369 NLRs annotated by R-Predictor.

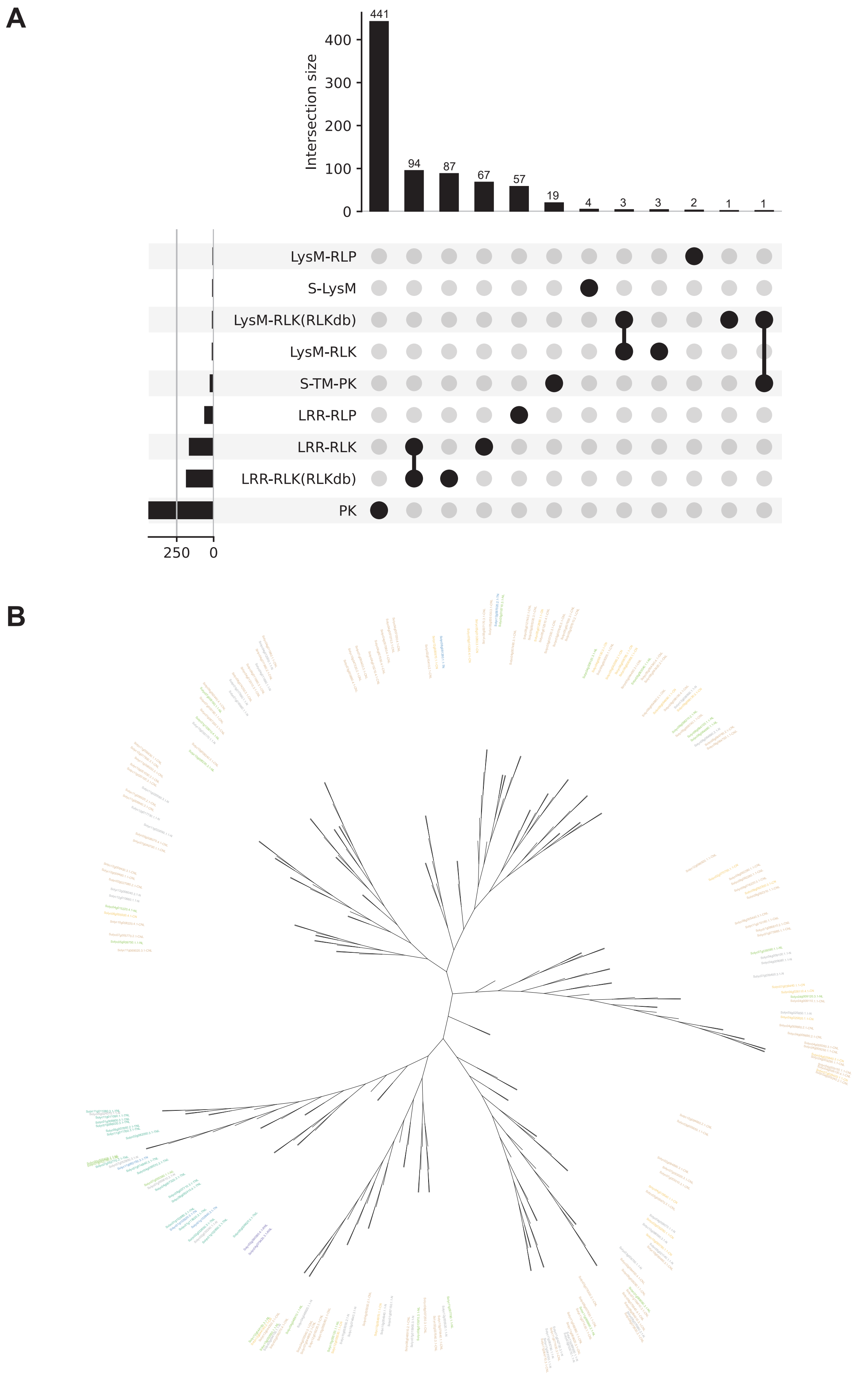


**Extended Data Fig. 6** | A. Upset plot of R-Predictor and RLKdb annotation results in ITAG4.0. B. A phylogenetic tree constructed based on 212 NLRs annotated by R-Predictor.

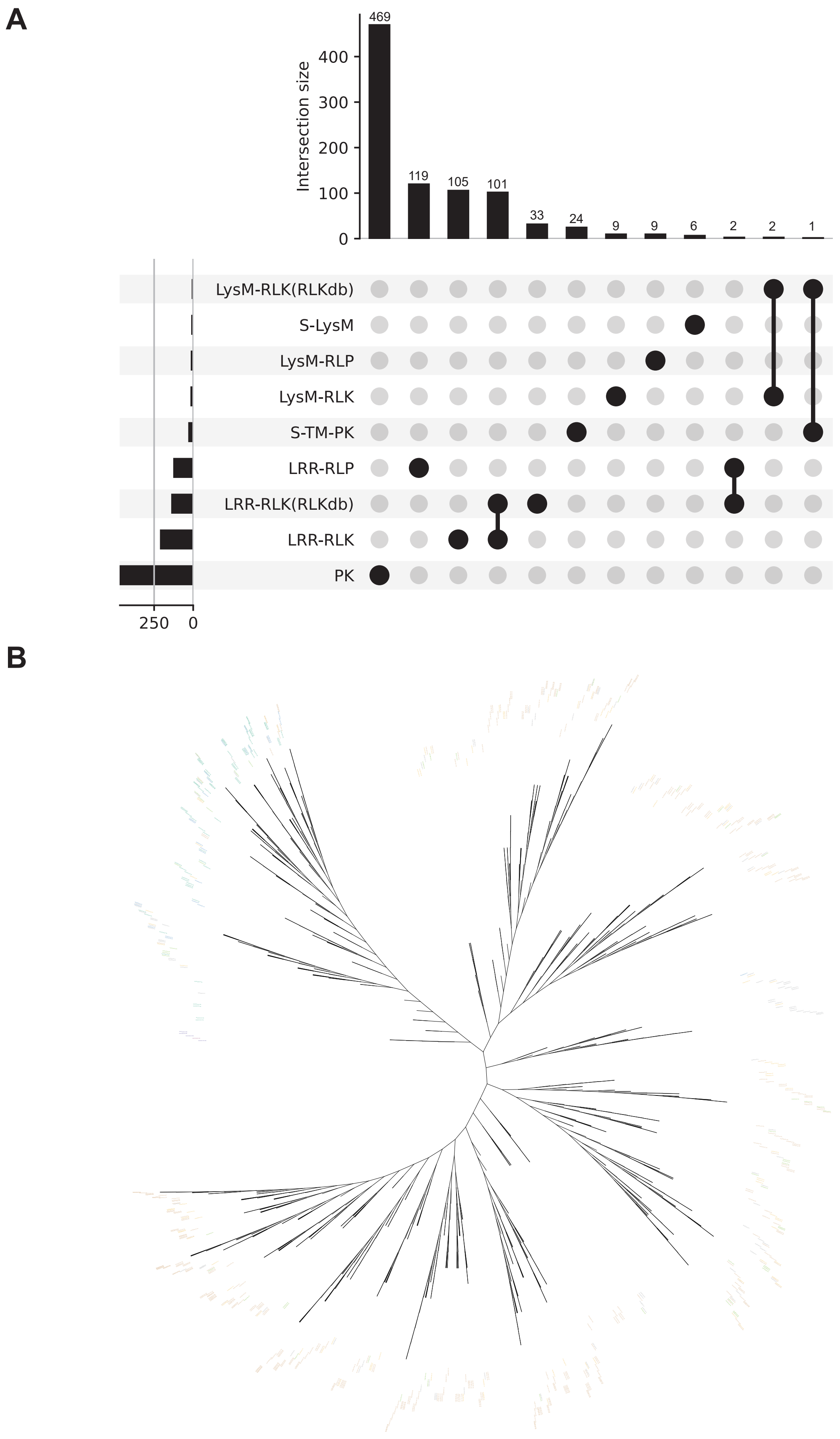


**Extended Data Fig. 7** | A. Upset plot of R-Predictor and RLKdb annotation results in PN40024_T2T. B. A phylogenetic tree constructed based on 745 NLRs annotated by R-Predictor.

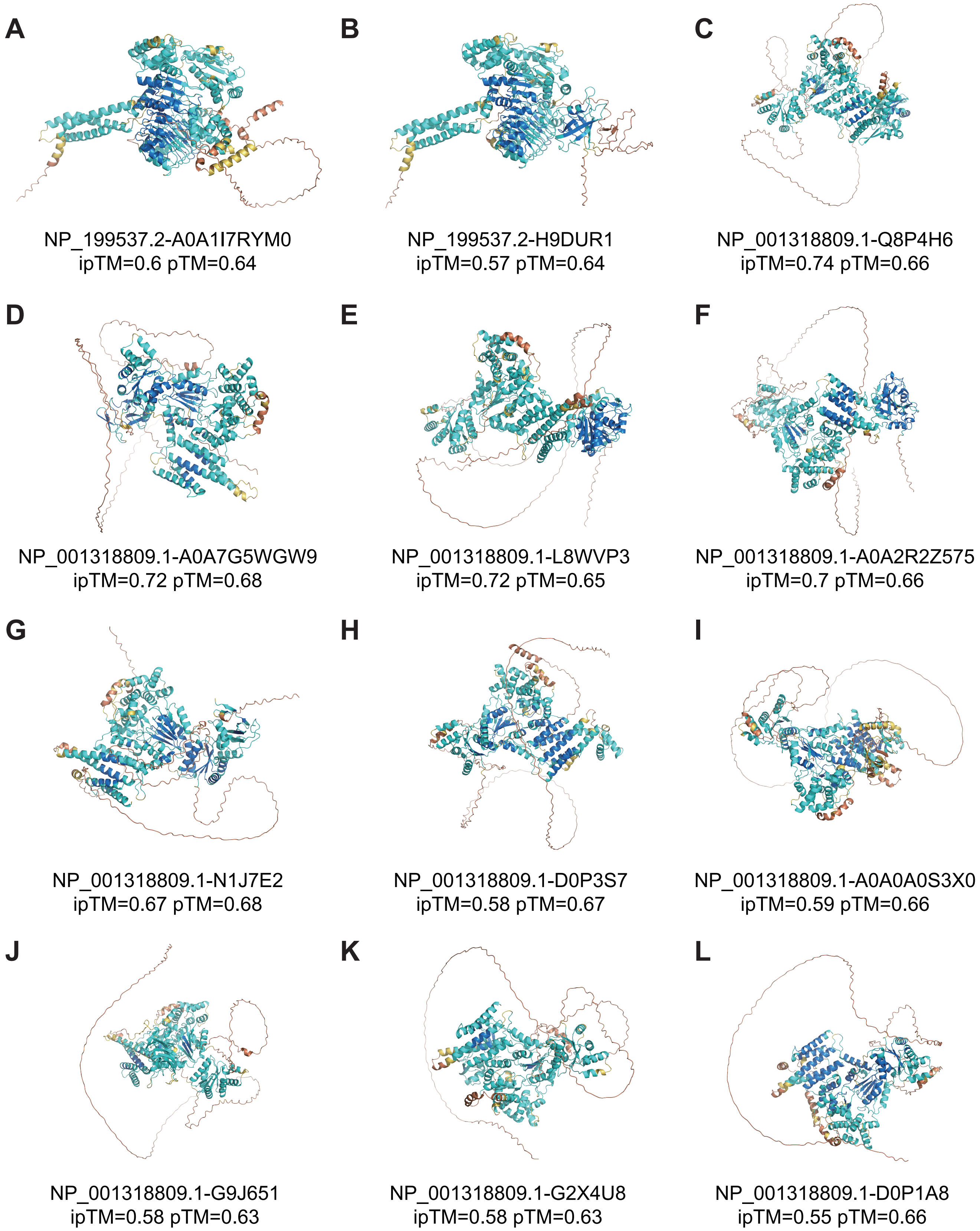


**Extended Data Fig. 8** | The three-dimensional structure of 12 putative NLR-effector complexes.

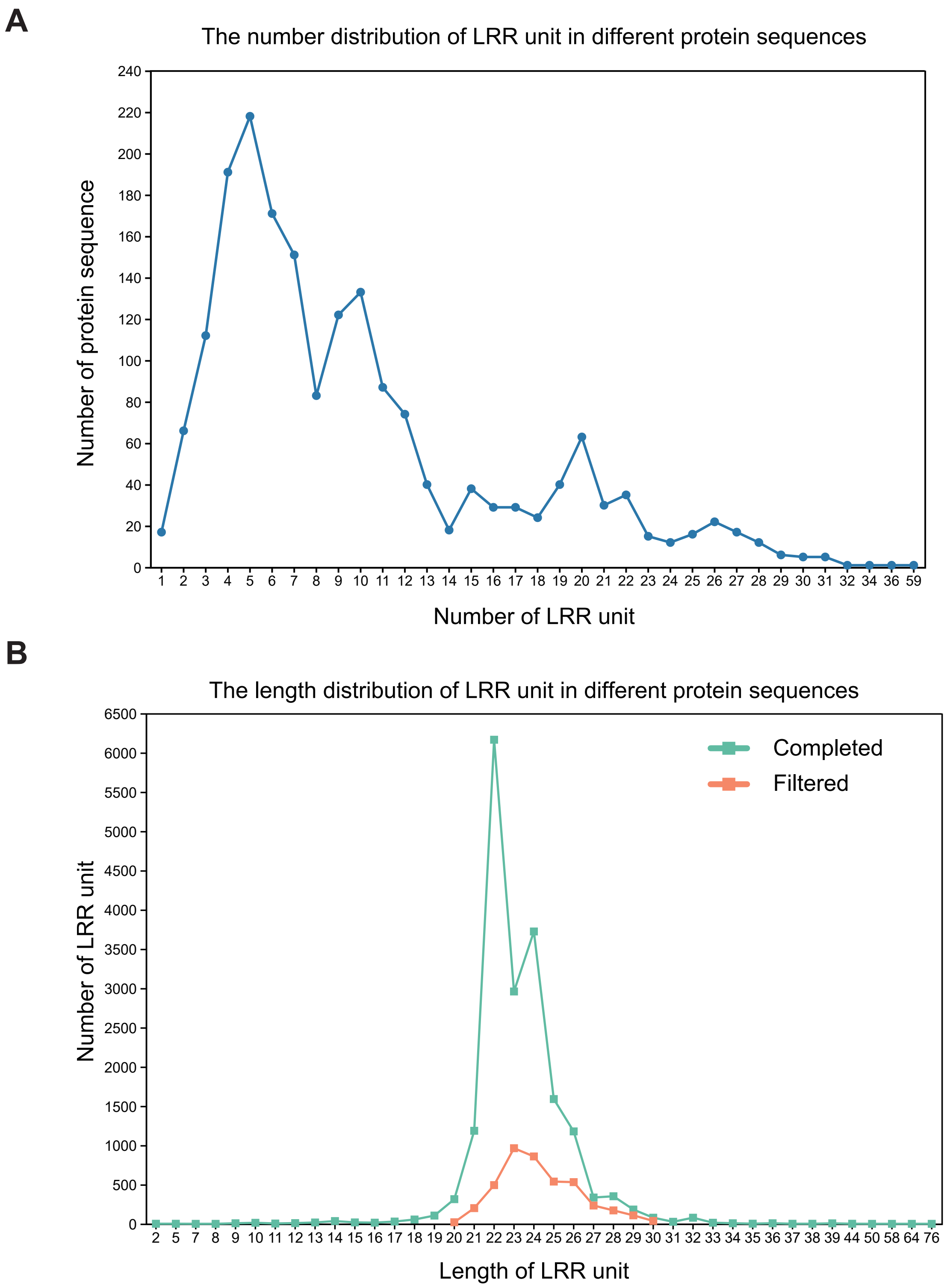


**Extended Data Fig. 9** | A. Number distribution of LRR units in 1,885 LRR proteins B. Length distribution of all LRR units of 1,885 LRR proteins. The light green line represents the original LRR unit data set without processing, and the orange curve represents the LRR unit data set after Z-score.
